# Inertial effects on work production in sub-maximally activated skeletal muscle

**DOI:** 10.64898/2026.05.01.722026

**Authors:** Colin Goodman, Brandon Reder, Lance Brooks, James Wakeling, Andrew A. Biewener, Nicolai Konow

## Abstract

Mass is a fundamental aspect of muscle contractile function, yet the inertial effects of inactive muscle mass is generally neglected in modeling and not quantified in studies on small muscles or isolated fibers. However, during submaximal contractions, inactive muscle tissue may take longer to be accelerated by active fibers, and may be subject to prolonged deceleration, both of which may potentially reduce force development and work output. We sought to test if inactive tissue mass imposes an inertial penalty on muscle performance, using *in situ* sinusoidal work-loop experiments on rat plantaris muscles. Regional fascicle dynamics, measured across supramaximal and submaximal levels of activation, showed that decreasing activation significantly reduced fascicle strain and increased both shortening and lengthening latency. Contrary to our predictions, however, reductions in work, beyond those explained by decreased fascicle strain, were negligible. Normalized work did not decline disproportionately relative to force, suggesting no clear inertial penalty on work at this muscle size. Our findings suggest that while inactive muscle mass influences the dynamics of submaximal contractions, its impact on work during submaximal contractions at small muscle sizes is limited.

## Introduction

Mass is fundamental to the function of a muscle’s contractile and elastic proteins and other cellular components. However, because most of what is known about whole muscle contractile behavior is based on single fiber or small muscle experiments, muscle mass is generally ignored in studies of the contractile mechanics of muscle (Mendoza et al., 2023). Additionally, muscle mass is typically incorporated as a component of segment mass in muscle models that drive musculoskeletal simulations (Anderson and Pandy, 2003; Chumanov et al., 2011; Delp et al., 2007; Hamner et al., 2010; John et al., 2013; Lai et al., 2021; Seth et al., 2018). Nevertheless, for submaximal contractions typical of most activities of daily living in animals including humans, inactive tissue mass must both be accelerated by actively recruited fibers that shorten to perform positive work and then be decelerated (Ross and Wakeling, 2016). Inactive muscle tissue, therefore, may exert an inertial effect that reduces the rate of muscle force development and the amount of work performed during submaximal contractions.

Evidence that inertial effects due to inactive muscle mass can impose a penalty on the speed of muscle shortening, the rate of force development, and the work produced by a muscle comes from both modeling (Günther et al., 2012; Ross and Wakeling, 2016; Ross and Wakeling, 2021) and experimental contractile studies on muscle (Holt et al., 2014; Ross et al., 2020). In these studies, penalties have been in the range of a 4-5% reduction in work for an 85-123% increase in effective mass (Ross et al., 2020) and a 2-to 2.5-fold reduction in maximal shortening velocity under submaximal activation conditions (Holt et al., 2014; Tijs et al., 2021). Modeling studies have also demonstrated that increased muscle mass reduces maximal shortening velocity and the rate of force development (Böl and Reese, 2008; Meier and Blickhan, 2000; Ross and Wakeling, 2016) of distributed mass Hill-type muscle models. In general, inertial effects are likely to be greater at larger body and muscle sizes, because muscle mass will increase at a greater rate than the scaling of muscle area and force. This is because while muscle mass scales with muscle volume, the force muscle can produce scales with the physiological cross-sectional area. Thus, for isometric scaling, the increase in inertial mass effects is predicted to scale α M^1/3^.However, it is unclear if inertial penalties influence the operation of muscles at the size-scale of the rat, a popular model of musculoskeletal function in mammals, including humans.

Based on the aforementioned studies, we hypothesize that inactive tissue mass significantly reduces the work that the actively recruited fibers of a muscle can produce during submaximal activations, which involve the recruitment of a fraction of the muscle’s fibers. To evaluate our hypothesis, we used *in situ* ergometry experiments carried out on the rat plantaris (PL). Contractile force and length recordings were made at both supramaximal (max) and submaximal (submax) activation of the muscle during sinusoidal work-loop experiments (Josephson, 1985) to isolate activation-dependent effects on muscle work. In addition, we recorded fascicle length changes at proximal, distal, and proximo-to-distal regions of the muscle belly using sonomicrometry (Eng et al., 2019;Taylor-Burt et al., 2020; Tijs et al., 2021) relative to overall muscle length-changes imposed by the motor lever.

These experiments tested the following hypotheses:

[HYP 1] Compared with supramaximal contractions, submaximal contractions undergoing sinusoidal length oscillations and activated during shortening incur an inertial penalty on the work performed by active fascicles that must accelerate the mass of inactive fibers when shortening. This will also result in slower rates of force development. As a result, the ratio of work performed during submaximal versus supramaximal activation will be less than the ratio of isometric tetanic force generated for each condition.

[HYP 2] The length changes or strain behavior of the muscle’s fascicles will diverge from the overall sinusoidal length changes of the muscle (imposed by the motor lever) due to inertial effects of inactive tissue mass and series elasticity within the muscle. Specifically,

[2a] Due to the accelerated mass of inactive fascicles as the muscle shortens, muscle fascicles will continue to shorten despite initial overall lengthening of the muscle by the motor lever, leading to fascicle-level work enhancement; whereas,

[2b] When the muscle is being lengthened by the motor lever, inactive fascicles will cause activated fascicles to shorten less rapidly at the onset of shortening, reducing the amount of work performed.

[HYP 3] Compared to supramaximal contractions, submaximal contractions while be characterized by reduced fascicle strains, resulting from decreased acceleration and velocity at lower activations.

## Methods

### Animals and study muscle

All experiments were carried out at the Concord Field Station, Bedford MA. USA in accordance with Institutional Animal Care and Use regulations of Harvard University (FAS; protocol 20_09-04). Experiments were conducted on plantaris (PL) muscles of fully-grown Sprague Dawley rats (*Rattus norvegicus*; *N* = 6, with an age-range of five to 12 months, and a body mass of 590 ± 170 g; Charles River, Wilmington, MA; Table 1). Animals of this large size were selected to provide plantaris muscles of a sufficient size for the experiment.

**Table 1.** Subject and plantaris muscle morphometrics.

| **Variable** | **mean** | **S.D.** |
| --- | --- | --- |
| Rat body mass (kg) | 0.59 | 0.17 |
| Muscle mass (g) | 0.64 | 0.25 |
| Muscle length<sub>measured</sub> (mm) | 43.80 | 3.78 |
| Muscle length<sub>interpolated</sub><sup>#</sup> (mm) | 43.90 | 3.80 |
| Fascicle length (mm) * | 18.94 | 1.46 |
| Pennation angle (°) * | 9.60 | 0.82 |
| PCSA (cm<sup>2</sup>) | 0.31 | 0.10 |

\*Denotes values derived from a separate representative sample of dissection. <sup>#</sup>Regression of animal mass against PL muscle length in cadaver dissections.

We chose the rat plantaris (PL) for study as it is well characterized and has been used in several relevant studies (Eng et al., 2019; Holt et al., 2014; Ross et al., 2020). The muscle contains a mix of primarily fast-contracting fiber types, including 11 to 63% type IIa, 31 to 38% IIx, and 46 to 47% IIb, as well as 5 to 9% type I fibers (Armstrong and Phelps, 1984; Caiozzo et al., 1992; Delp and Duan, 1996; Eng et al., 2008). The muscle architecture is near parallel-fibered with fiber resting angles of 9.6 ± 0.8° (Table 1).

### Instrumentation procedure

Animals were anaesthetized on isoflurane (Induction in a chamber; 4%, 2L/min O_2_, followed by maintenance on a mask; 2–3%, 0.8L/min O_2_). Since the procedures often lasted 6-7 hours, normal saline (2ml) was administered subcutaneously, and the bolus was replenished when resorbed. Core temperature was maintained at 38.5–39.5 °C using a heat-pad, supplemented with infrared light, and monitored using a rectally inserted thermocouple.

To facilitate whole-field tetanic stimulation (supramax) and stimulation at reduced activation levels (submax) across the entire muscle belly, a custom-made bipolar nerve cuff (silver wire, interpole spacing of 8 mm) was securely fitted around the sciatic nerve as close as possible to the nerve’s exit from the spinal cord. The nerve was cut proximally to the cuff to avoid retrograde stimulation of the CNS and sutured to the underlying musculature. The proximal placement of the cuff was to ensure that stimulation levels did not change due to cuff movement over the nerve during work loop contractions. The skin was cut around the ankle and gently retracted to a level above the knee to facilitate isolation of PL via blunt dissection for insertion of sonomicrometry crystals. The tendons of gastrocnemius, soleus, tibialis anterior, and the extensor digitorum complex were cut and the muscles reflected and resected, leaving only plantaris and the small digit flexors intact in the posterior compartment. The tendon of plantaris was then cut at a level distal to the calcaneus and 1-0 silk suture was tied to the free tendon at the myotendinous junction (to reduce series compliance) using a sliding knot, which was enforced with cyanoacrylate gel.

To measure muscle length-change independently of the servomotor oscillations during the experiment, three sonomicrometry crystals (1.0mm diameter, Sonometrics Corp) were implanted along the muscle belly axis, one at the distal-most position possible, tracking the myotendinous junction, one at the proximal-most position that avoided interference with the muscle’s nerve-supply, and one intermediate to the other two crystals. This approach resulted in measurements of segment distances of approximately 6–12 mm (distal-mid, mid-proximal) and 12–24 mm (distal-proximal) respectively (Figure 1). Sonomicrometry signals were monitored on an oscilloscope and tuned, based on trigger level relative to receiving signal strength, as the lead wires were secured using a suture of 5-0 silk through the epimysium approximately 3-5 mm from the crystal, in order to minimize motion artifacts such as level-shifts during work-loops.

**Figure 1.**
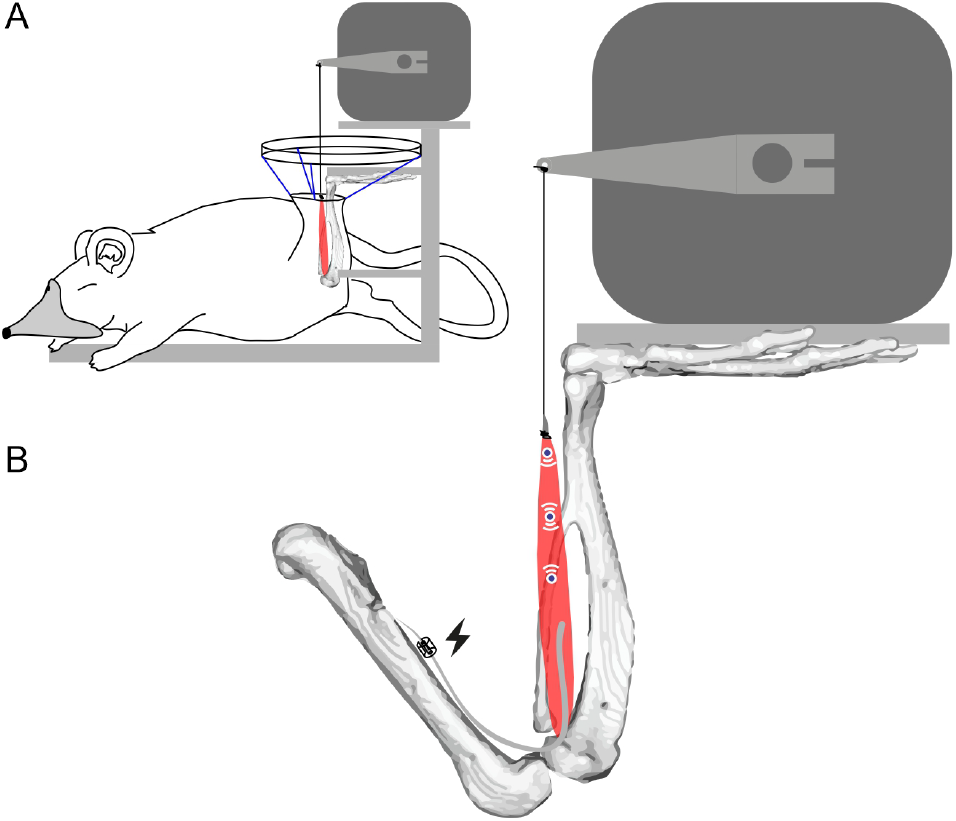
Experimental setup. (A) Subject arranged in stereotaxic frame (grey) holding servomotor and clamping left foot and knee, with shank skin tented up using elastics (blue) to form a bath filled with mineral oil. (B) close-up of muscle preparation with nerve cuff 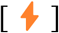 on proximal sciatic nerve and sonomicrometry implants [((O))] in PL muscle belly (red).

The rat was then placed prone on a heat pad in a custom-built stereotaxic rig, with the knee clamped at approximately 90° and the foot secured to an acrylic foot plate with the ankle at approximately 90° (Figure 1). The tendon was tied to the lever-arm of a servomotor (305B-LR Aurora Scientific Inc. Aurora ON) and the nerve cuff was connected to a stimulator (S48 Grass Technologies, West Warwick RI). The loose skin of the shank was carefully pulled back to re-envelop the lower limb, and tented up to the stereotaxic frame using skin clips and rubber bands. This formed a bath around the muscle preparation that was filled with warmed (38°C) mineral oil (Holt et al., 2014; Tijs et al., 2020). The bath and muscle were kept at a temperature comparable to the animal’s core (38 ± 1.5°C) using an infrared lamp and continuously monitored via a thermocouple signal that was recorded to IgorPro throughout the experiment.

### Muscle preparation conditioning

After rigging, and prior to any stimulation, preparations were acclimated for 15 minutes. Then, a series of tetanic contractions were elicited, with the muscle at a passive length roughly corresponding to *L*_0_, to remove any compliance within the preparation. Preparations were allowed three minutes of rest between all tetanic contractions to minimize impacts of fatigue. Once post-tetanic passive force had stabilized, a series of twitches were elicited at increasing voltages to determine the supramaximal stimulation voltage (typically 2–8V).

### Force-Length measurements

To perform our main experiment, we first characterized the force-length relationship for each muscle preparation. A series of tetanic stimuli across increasing muscle lengths were used to measure the force-length curve and determine peak tetanic force (*P*_o_) and muscle length at peak tetanic force (*L*_o_) for the preparation. The proximal origin of PL on the femur made it impossible to reliably measure total muscle length during the experiment. Therefore, prior to the study, we measured muscle length and body mass in cadavers (*N* = 5) and used a regression to estimate resting length (*L*_r_) for each experimental preparation, based on subject mass. Interpolated lengths fell very close to our *ex vivo* measurements (*R*^2^ = 0.90, *F*_1,4_, *p*<0.01; Table 1).

### Work loop approach and experimental conditions

The main goal of our study was to determine how inactive muscle mass influences muscle mechanical work. To this end, we used the work loop technique (Josephson, 1985) with cyclic, sinusoidal length changes imposed using the servomotor in concert with timed stimulation volleys. We chose input parameters aimed at generating maximal positive work during the shortening portion of each contraction cycle, specifically by 1) pre-stimulating the muscle by -5% of the shortening phase to reach high force before muscle shortening; 2) discontinuing stimulation at a duty cycle of 0.45 to minimize active force production during muscle re-lengthening (Baird et al., 2021; Swoap et al., 1997); 3) centering the muscle strain trajectory around *L*_o_ and keeping strain trajectory low (±3% *L*_o_) (Shelley et al., 2024); and 4) altering stimulation frequency to ensure fusion of tetani even at submaximal activation (85-125Hz)(Shelley et al., 2022). Whist selecting our parameters, we also sought to facilitate comparisons of our results to those from prior simulation studies (Ross et al., 2018). Both stimulation pulse width (1 ms) and oscillation frequency (4Hz) were kept constant for all experiments. Representative cycles of example work loop patterns, along with two sets (4 cycles each) of maximal and submaximal work loops are shown in Figure 2.

**Figure 2.**
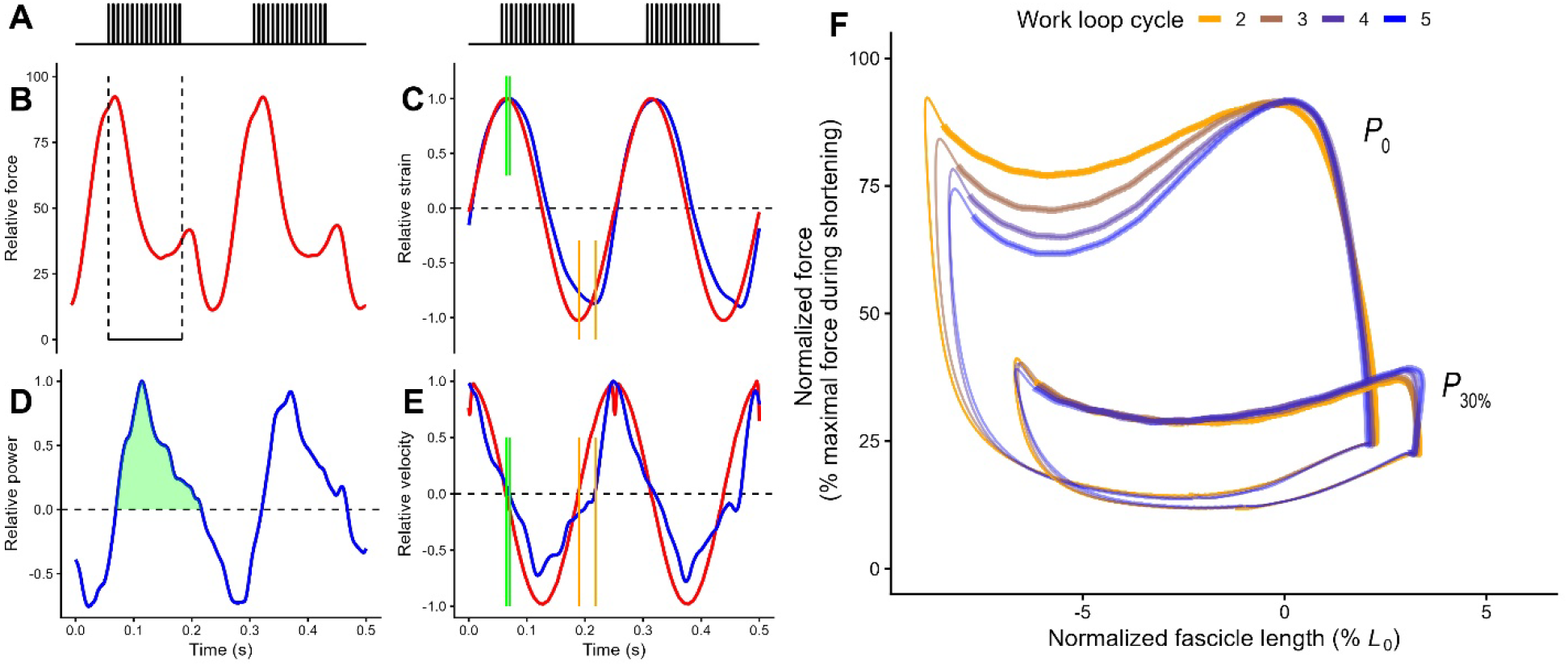
Sample work loop trials showing A) muscle stimulation, B) force (relative to *P*_o_), C) relative displacement of the motor lever (red) versus proximal fascicle segment (blue), D) relative power of the proximal fascicle segment and E) relative velocity of the lever (red) versus the proximal fascicle segment for two cycles over time. B) Shortening cycling force denoted by vertical dashed lines was determined by the average force during the period of motor shortening under the time-varying force curve during stimulation. D) Positive cycle work denoted by the green shaded region was determined by the positive area under the power curve. E) Shortening and lengthening latency is represented by the two pairs of black vertical lines, defined as the latency between the transitions in motor lever lengthening to shortening versus fascicle lengthening to shortening; and the transition in motor lever shortening to lengthening and fascicle shortening to lengthening (zero velocity). Note that for ease of interpretation in B-E, x-axis values were normalized against their respective maxima. F) Representative work loops at maximal (*P*_0_) and submaximal (*P*_30_) stimulation levels. Fascicle length is shown as a percentage of optimal fascicle length (*L*_0_) at *P*_0_. Force is shown as maximum force during maximum fixed-end tetanic stimulation at *L*_0_. Colors delineate consecutive cycles and thick line segments indicate the period of stimulation. Note that the first work loop cycle is not used in the analysis and thus not plotted.

To elicit submaximal force and work production during cyclic sinusoidal contractions, we systematically varied stimulation voltage. Prior work has demonstrated that decreasing the stimulation voltage results in activation of decreasing volume of a muscle(Holt et al., 2014; Tijs et al., 2021), in line with our experimental goal. Given the multiple variables that potentially could affect muscle work (e.g., temperature fluctuations, fatigue or damage, and movement of the nerve cuff), we designed the experiment to ensure that tetanic force production and work production at a given activation level were as closely timed as possible. We lowered voltage to approximately 30% of the supramaximal level and used brief tetani to determine if force had dropped to ∼30% of that achieved at supramaximal activation. We then collected a 250 ms fixed-end tetanus and verified that the muscle could sustain a plateau (*P*_30%_), followed by the work-loop trial (*N* = 5 cycles). Stimulation voltage was then systematically increased, and the protocol was repeated (∼45%, ∼70%), finishing with paired fixed-end tetanus and work loop trial pairs at supramaximal stimulation.

All experiments ended with a final fixed-end tetanus at *L*_0_ to determine if the preparations had suffered damage or fatigue during the experiment, by the 10% drop criterion. Apart from one preparation that experienced a decline in force of 17%, all preparations fell below this criterion. The inclusion of this individual did not alter the results of this study. When an experiment was concluded, the deeply anesthetized rat was euthanized using an overdose of sodium pentobarbital (IC), followed by thoracotomy and heart removal. The plantaris muscle was carefully dissected free and the relaxed length of the muscle belly (*L*_R_) as well as the sonomicrometry segments were measured to the nearest 0.1 mm. The sonomicrometry crystals were removed and, after cutting away both tendons, the muscle was weighed to the nearest 0.01 g.

### Data acquisition, measurement post-processing and summary data extraction

Data on stimulation intensity and duration, muscle fascicle length, and muscle lever force and length were acquired by a PC running IgorPro (Wavemetrics, Lake Oswego, OR) via a NIDAQ (USB-6212). Analogue signals were sampled at 10 kHz. Prior to data extraction, sonomicrometry signals were visually inspected for level-shifts (manifested as artifactual jumps in fascicle length). When level-shifts were small in duration, length values were removed to facilitate accurate interpolation. Level-shifts that were greater in duration were corrected using a custom Igor script. Any signals wherein level-shifts could not be unambiguously identified and corrected were not used for data extraction or subsequent analysis. This criterion resulted in retention for analyses of a total of 13 fascicle segments (proximal: *n* = 5; distal: *n* = 5; whole muscle segment: *n* = 3)

All raw measurements were conditioned using a quintic smoothing spline, and the level of conditioning was monitored visually and carefully adjusted for each signal to avoid over- or under-smoothing. Using the conditioned measurements, a custom IgorPro script was used to calculate the time-varying velocity and power of the lever motor, and the muscle fascicle segments. Velocity was calculated as the first derivative of length, while power was calculated as the product of lever motor force, and lever motor or muscle fascicle velocity. From the conditioned and processed signals, the following variables were extracted for further analysis: maximal isometric force (*P*_0_), optimal length (*L*_0_), average cycling force during shortening, positive cycle work, strain, and cycling onset and offset latency. *P*_0_ was defined as the maximal difference between active and passive force during isometric tetani and *L*_0_ was the whole muscle length at which *P*_0_ occurred. Positive cycle work was defined as the positive area under the time-varying power curve. Although this typically corresponded to the shortening cycle of the motor, fascicle segments often continued shortening for a short period of time while the motor lever was lengthening.

To test our hypothesis that inactive muscle mass imposes an inertial penalty during submaximal contractions, we used a suite of generalized linear mixed models (GLMM). Because neither consecutive work loop cycles nor fascicle segments within a preparation are independent, we averaged the values for each response variable and predictor across cycles and fascicle segments. Additionally, because of the ubiquity of length-history effects on contractile properties (Hahn et al., 2023; Abbott and Aubert, 1952; Herzog, 1998; Edman et al., 1982), we discarded the first full work loop cycle in each trial, and only included cycles 2-4 in calculations of averages. This resulted in a total of 22 trials from six animals retained for analysis.

First, to test our prediction that total strain would decrease with decreasing stimulation, we used a GLMM with normalized strain (expressed as a percentage of *L*_0_) as the response variable, normalized force (expressed as a ratio of *P*_0_ at *L*_0_) as the response variable, and preparation (animal ID) as a random effect. Additionally, because of the potentially disparate effects of activation level on fascicle shortening vs lengthening, we also ran separate models for normalized shortening and lengthening strain. We predicted that if inactive muscle mass imposes an inertial penalty during submaximal contractions, then the positive work performed by the muscle should be underpredicted by the force produced by the muscle, resulting in a slope greater than one between relative force and relative work. To test this, we used a GLMM with the ratio of maximal positive work (during maximal stimulation) as the response variable, normalized force (during maximal stimulation) as the predictor, and individual as a random effect. Since inertia may cause a change in fascicle shortening velocity, thus limiting fascicle excursion for a given oscillation frequency, we included normalized strain as a covariate. Additionally, to test whether the coefficient of the positive work ratio regressed against the isometric force ratio differed significantly from one, we subtracted the ratio of maximal isometric force from the ratio of maximal positive work.

We also predicted that an inertial penalty would result in greater shortening as well as lengthening latencies during submaximal stimulation. To test these predictions in kind, we used two separate GLMMs. One tested for effects of stimulation level on shortening latency by using shortening latency (expressed relative to latency at maximal stimulation) as response variable, normalized force as the predictor, normalized shortening strain as a covariate, and individual as a random effect. The second model tested for effects of stimulation level on lengthening latency, using lengthening latency as the response variable, normalized force as the predictor, normalized lengthening strain as a covariate, and individual as a random effect. For all models, effects were considered significant if *p*<0.05.

## Results

Muscle preparations included for analyses (*N* = 6) developed specific tension of 14.0 ± 7.6 N/cm^2^ at tetanic *L*_0_, which was 46.4 ± 4.4 mm across the six preparations. Normalized positive work increased with both stimulation level and strain, with all but one submaximal stimulation trial falling below the line of isometry with respect to normalized work-loop force (Figure 3; Table 2). However, the regression slope between normalized force and positive work was not significantly greater than one (*t*_14_ = 0.5, *p* = 0.62; Figure 3; Table 2). The effect of strain on normalized positive work was not significant after controlling for stimulation level (*t*_14_ = 0.34, *p* = 0.74; Table 2).

**Figure 3.**
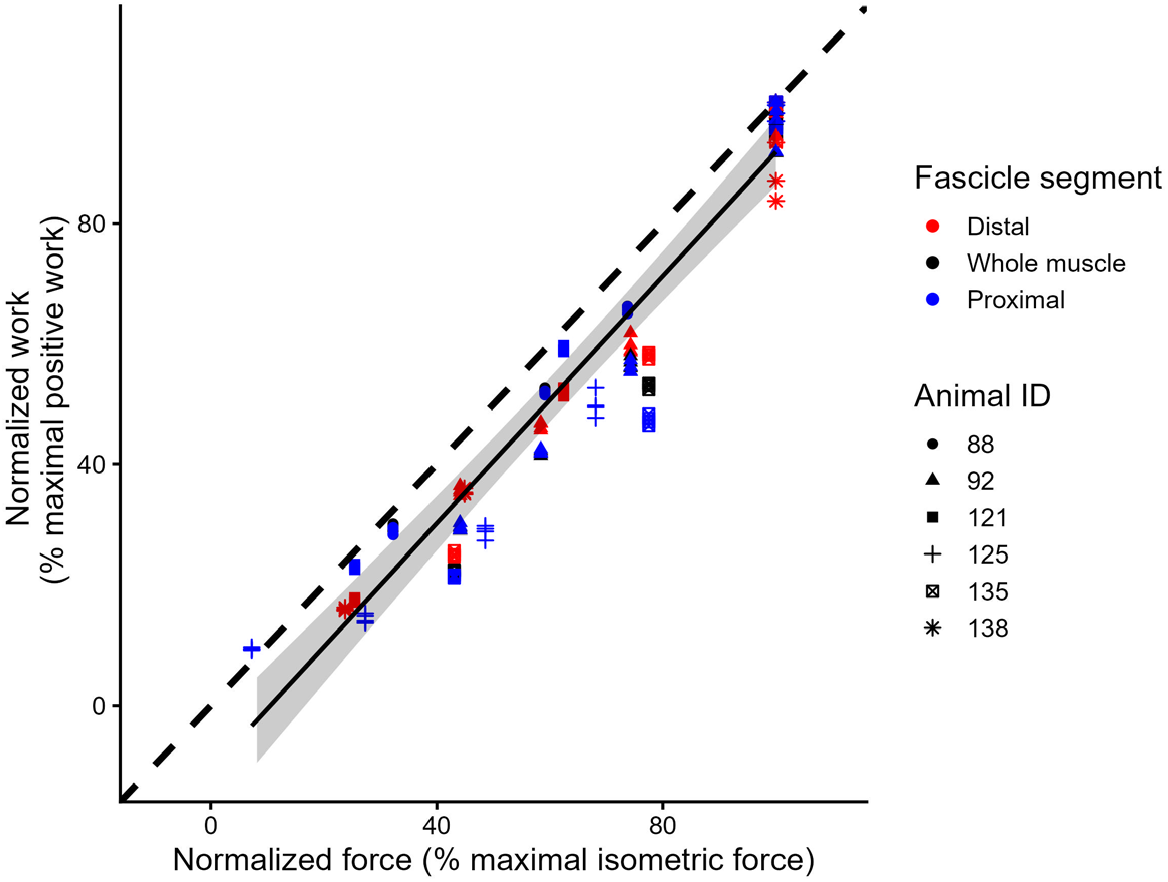
Relationship between stimulation-normalized work and isometric force. Force values were normalized as in Figure 2F. Positive work values were normalized to the cycle that produced maximum positive work during maximal stimulation. Colors denote fascicle segments, and shapes denote muscle preparations. The solid black line represents the conditional relationship between normalized force and work, holding strain constant. The shaded gray region represents the 95% confidence interval for this relationship. The dashed line represents the line of isometry (i.e. no inertial penalty); values falling below this line indicate that normalized work is less than expected for a given normalized force. Note that differing cycles and fascicle segments are represented on the graph but were pooled for the analysis.

**Table 2.** Summary statistics from generalized linear mixed models.

| Model | Variable | Slope | S.E.M | d.f. | t | p |
| --- | --- | --- | --- | --- | --- | --- |
| Total strain ~ normalized force | Normalized force | 3.9 | 0.49 | 15 | 7.97 | **<0.001** |
| Shortening strain ~ normalized force | Normalized force | -4.8 | 0.52 | 15 | -9.2 | **<0.001** |
| Lengthening strain ~ normalized force | Normalized force | -0.89 | 0.46 | 15 | -1.91 | 0.07 |
| (Normalized work - normalized force)~ normalized force + total strain | Normalized force | 0.03 | 0.06 | 14 | 0.5 | 0.62 |
|  | Total strain | 0.002 | 0.006 | 14 | 0.34 | 0.74 |
| Lengthening latency ~ normalized force + lengthening strain | Normalized force | -8.32 | 2.71 | 14 | -3.07 | **<0.01** |
|  | Lengthening strain | -0.34 | 0.96 | 14 | -0.35 | 0.73 |
| Shortening latency ~ normalized force + Shortening strain | Normalized force | -5.81 | 1.39 | 14 | -4.19 | **<0.001** |
|  | Shortening strain | -0.096 | 0.183 | 14 | -0.53 | 0.61 |

All models included subject as a random effect. Boldfaced p-values represent statistically significant effects.

Patterns of fascicle strain differed significantly across stimulation levels, with increased stimulation resulting in greater total strains (*t*_15_ = 7.97; *p* <0.001; Figure 4; Table 2). This pattern was largely driven by fascicle strain during shortening (*t*_15_ = -9.2; *p* <0.001; Figure 4; Table 2), as lower stimulation resulted in greater strain during lengthening (*t*_15_ = -1.91; *p* = 0.07; Figure 4; Table 2). Additionally, after accounting for the differences in strain (expressed as percent maximal strain), most of the normalized work values fell above the line of isometry. This shift suggests that most of the reduction of normalized positive work at a given stimulation level may be explained by reductions in total strain (Figure 5).

**Figure 4.**
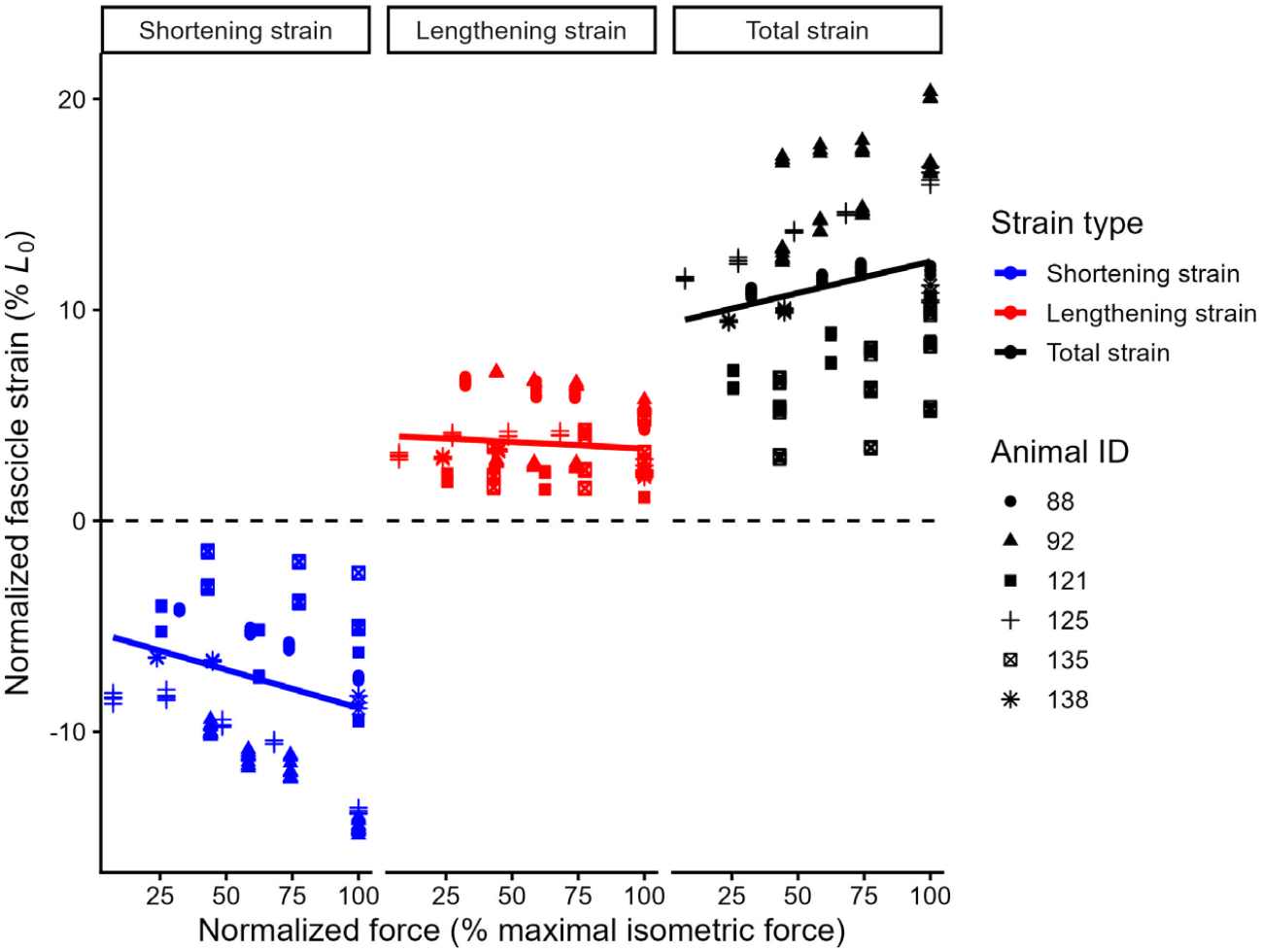
Changes in fascicle contractile strain with increasing activation. Force values on the x-axes are normalized to maximal isometric force during maximal stimulation, for each muscle preparation. Strain values on the y-axes are normalized against L_0_ (horizontal dashed line) for each preparation. Shapes denote preparations. Colors denote strain during shortening (blue; left panel), strain during lengthening (red; middle panel), and total strain (black; right panel). Decreasing values of shortening strain indicate increased shortening, while decreasing values of lengthening strain indicate decreased lengthening.

**Figure 5.**
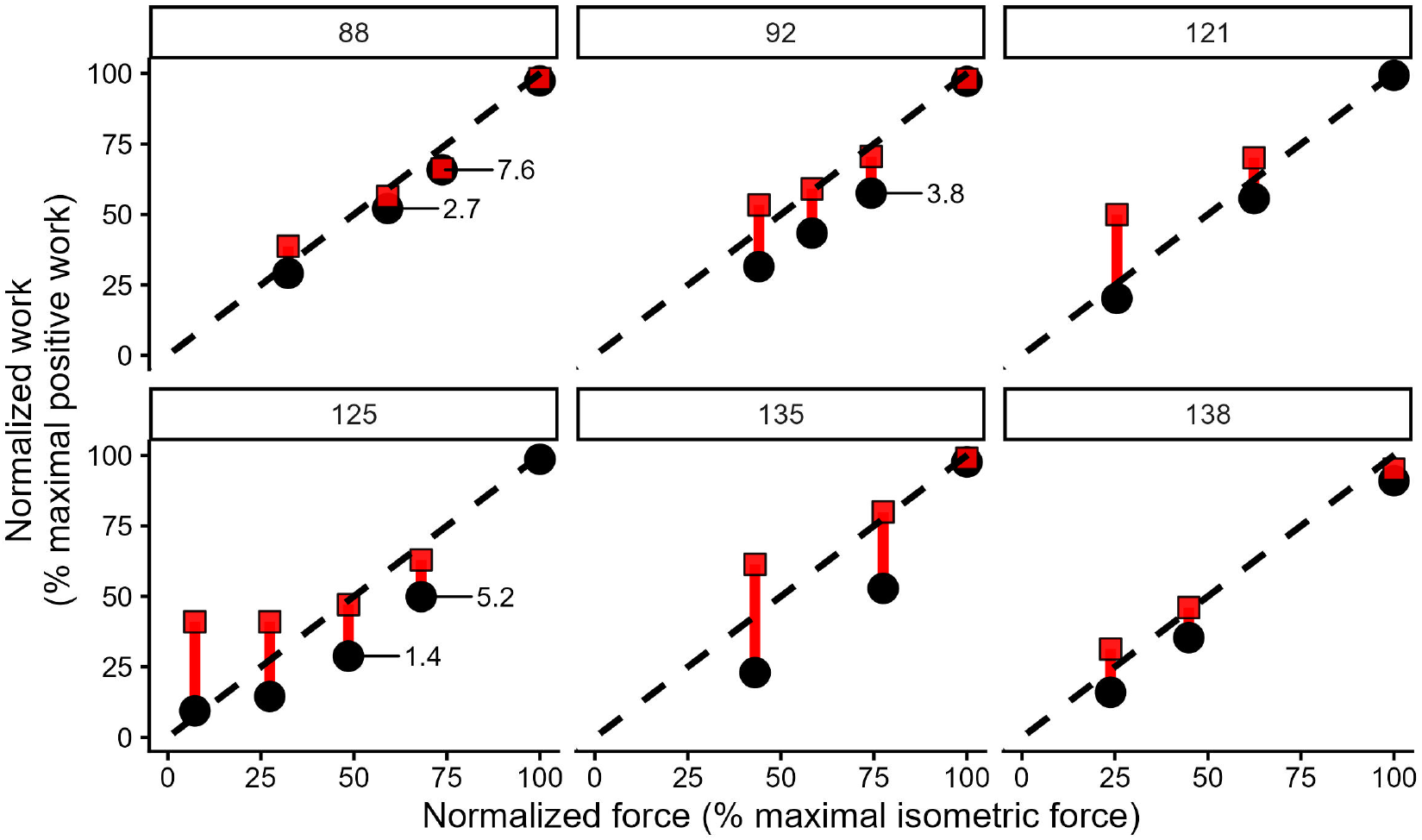
Residual-corrected work output. Data from Figure 3, shown with correction fascicle excursion. Larger, black dots represent empirical values, averaged across cycles and fascicle segments. Red squares represent the raw values, plus the percentage decrease in excursion during each submaximal stimulation. Work-force residuals were calculated as the distance between each point and the line of isometry. Red lines that cross the line of isometry are cases where the work-force residuals were smaller than the change in excursion from max stimulation. Red lines that do not reach the line of isometry are cases where work-force residuals were greater than the change in excursion from max stimulation, the magnitude of which is denoted using numeric labels. Red squares that fall below the line of isometry indicate that even after correcting for excursion, work remains less than predicted from stimulation level.

Patterns of fascicle contractile latency differed significantly across stimulation levels. We found an increase with lower stimulation of both the shortening latency (being the delay at which fascicles reached peak length after the motor, orange colored vertical lines in Figure 2C) and the lengthening latency (being the continuation of fascicle shortening into the period where the muscle was stretched by the motor, green colored vertical lines in Figure 2C), even when controlling for the decrease in fascicle strain. The effect of stimulation level on fascicle contractile latency was greater for lengthening (*t*_14_ = -3.07; *p* <0.01; Figure 6; Table 2) than for shortening (*t*_14_ = -4.19; *p* <0.001; Figure 6; Table 2).

**Figure 6.**
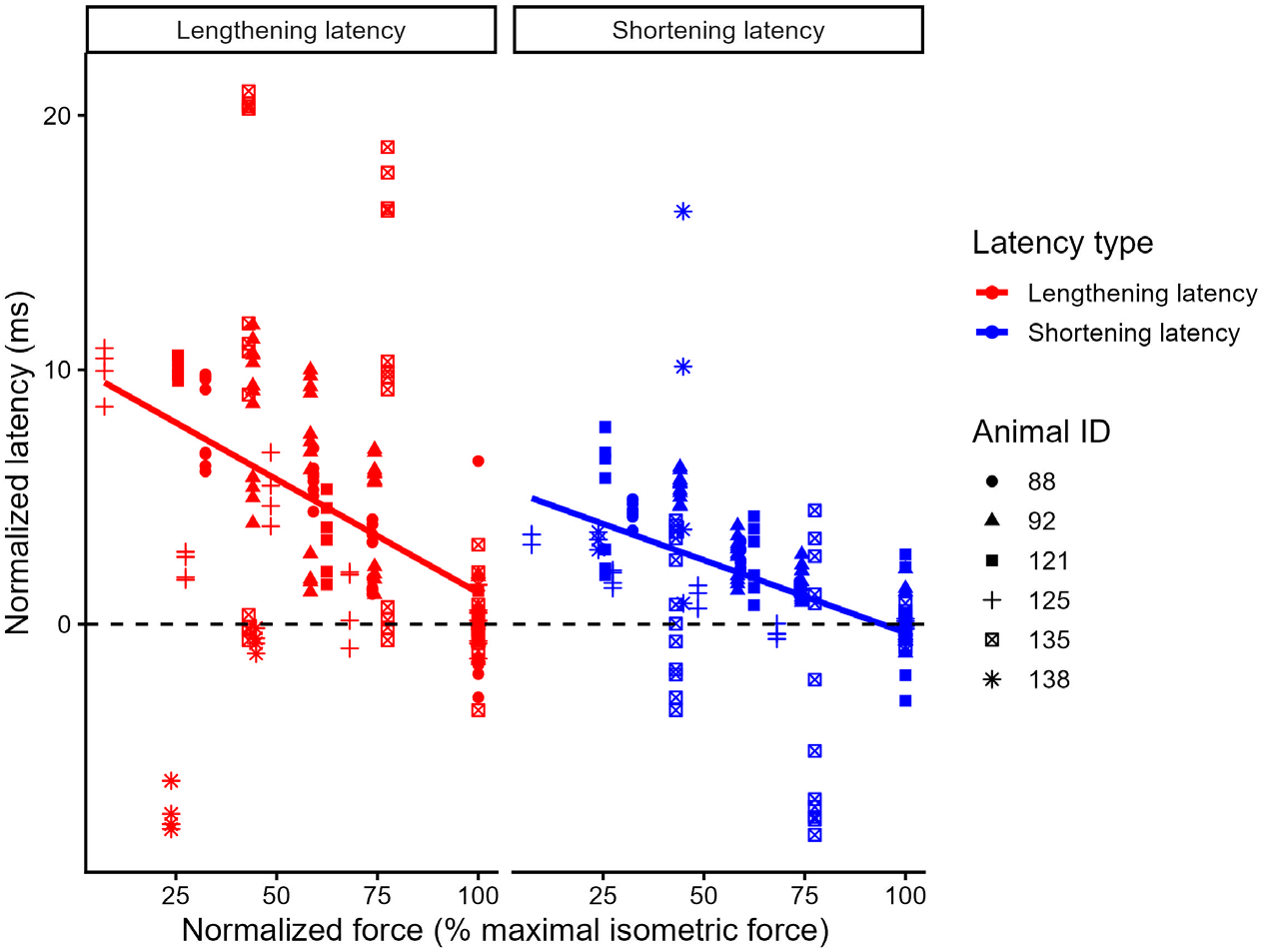
Differences in latency of fascicle shortening and lengthening across stimulation levels. X-axis values represent stimulation-normalized isometric force as described in Figure 4. Y-axis values represent the latency between lever transitions from shortening to lengthening (red) and lengthening to shortening (black). Latency values were normalized against the average latency at maximal stimulation for each muscle preparation. Positive values at a given stimulation level indicate that fascicle lengthening occurs later than at maximal stimulation. Negative values indicate fascicle lengthening occurs earlier than at maximal stimulation.

## Discussion

Maximal performance is costly to the underlying muscles that power it, and as a result, many organisms rely on submaximal muscle performance for routine activities. However, most studies that have investigated isolated muscle performance have done so with the muscle maximally activated, which excludes a critical component of muscle function. We sought to determine the impacts of inactive tissue mass on muscle undergoing submaximal contractions. To accomplish this, we measured time-varying force and strain of the rat plantaris muscle tendon unit (motor) and fascicles during sinusoidal work-loops at multiple activation levels. We predicted that inactive mass resulting from submaximal muscle stimulation would cause a disproportionate reduction in the work performed by the muscle, muscle fascicle strain magnitude, and the latency between movements of the motor and contractile excursions of the muscle fascicles. We found mixed support for these predictions: Although the positive work done by the muscle was nominally lower than expected for a given stimulation level (Figure 3), this effect did not withstand statistical testing across subjects, muscle regions, and contractile events. Further, we found that for most of our work loop trials, reductions in work at lower stimulation levels relative to supramaximal muscle stimulation could well be explained by associated reductions in fascicle strain (Figure 6). Consistent with our predictions, we found that total fascicle strain was significantly reduced with decreasing activation (Figure 4; Table 2). Finally, consistent with our predictions, we found that lower levels of stimulation resulted in modest fascicle lengthening and shortening latencies relative to length changes of the whole muscle, as imposed by the lever motor (Figure 6).

### Work during submaximal contractions

We did not find consistent evidence that relative positive work generated by the rat plantaris decreased with reduced stimulation level. However, both the magnitude and direction of our results are broadly consistent with results from an experiment that reported a 4.7% reduction in positive work when rat PL was supramaximally stimulated after its mass was artificially increased by 123% (Ross et al., 2020). Scaling the result of Ross et al., (2020) to our framework corresponds to an approximate 4.7% work reduction at approximately 45% activation; whereas our estimated regression slope of 1.03 (Figure 3) predicts only an approximate 1.6% work reduction at the same level of activation. Protocol differences likely explain some of this discrepancy, as Ross et al. (2020) varied strain amplitude (± 5–10% *L*_o_), thereby increasing fascicle shortening velocity and acceleration at a given frequency and potentially amplifying inertial penalties. By contrast, we used a constant, smaller strain (± 3% *L*_o_), which may have attenuated inertial effects. However, two features of the Ross et al (2020) study would presumably reduce inertial contributions as compared to our study: a lower oscillation frequency (2 Hz versus 4 Hz) and the use of lever motor excursion as a proxy for muscle strain (measured here using sonomicrometry). We found that fascicle shortening often exceeded lever excursion (Figure 4), indicating that motor excursion likely underestimated fascicle strain—and thus inertial effects—in the preparations of Ross et al. (2020). Indeed, the changes in fascicle strain we measured were sufficient to explain most if not all of the reduction in fascicle work at reduced stimulation levels (Figure 5). Taken together, this finding is consistent with the relatively small effects predicted by models that incorporate scaling approaches to determine the critical mass and size of skeletal muscles at which inertial effects would manifest on force production (Chen et al., 2026) and muscle coordination (Latreche et al., 2026).

### Effects on fascicle strain during submaximal contraction

Across levels of stimulation, we found significant differences in total fascicle strain, with model-predicted strains (based on the GLMM results reported in Table 2) of 12.9 and 15.6% *L*_0_ for *P*_30_ and *P*_0_, respectively (Figure 4; Table 2). Importantly, however, this strain depression was despite the fact that there were opposing patterns of lengthening and shortening fascicle strains (Figure 4; Table 2), with a decrease in model-predicted fascicle lengthening from 3.75 to 3.13% *L*_0_ and an increase in shortening from 9.12 to 12.5% *L*_0_ for *P*_30_ and *P*_0_, respectively. Because the muscle was not stimulated until near the end of the lengthening cycle, most of the lengthening strain was passive and caused by the lever motor. Indeed, by comparison to shortening strain, fascicle lengthening strain more closely matched motor strain at all stimulation intensities (Figure 4). As a result, the observed differences in fascicle lengthening strain inherently arose during the 5 ms of motor lengthening (immediately preceding motor shortening) when stimulation of the muscle began. In order to resist lengthening and (or) to shorten during motor lengthening, fascicles must rapidly generate force, and their capacity to do so is likely diminished at submaximal stimulation intensity (Holt et al., 2014; Tijs et al., 2021).

The effect of reduced strain during active shortening is likely caused by a reduction of fascicle shortening velocity and acceleration during submaximal stimulation. At submaximal stimulation, actively recruited fascicles must accelerate inactive tissue mass, thereby increasing effective muscle mass. With oscillation frequency being constant in our experiment, the increase in effective mass likely reduced the strain achieved by active fascicles over the shortening period. Thus, this effect would be expected even if shortening strain for a given relative load was invariant across stimulation levels. Studies using *in silico, in situ*, and *in vivo* approaches have confirmed this reduction in active shortening velocity both at the level of the muscle tendon unit and its fascicles (Ross and Wakeling, 2016; Ross et al., 2018; Holt and Azizi, 2016; Tijs et al., 2021). One way to determine if the increase of effective mass drives this reduction in fascicle shortening strain would be to alter motor oscillation frequency. If oscillation frequency were increased, one would expect a reduction in fascicle shortening strain at both submaximal and supramaximal stimulation. However, given potential inertial effects, increasing oscillation frequency should have a greater effect on fascicle shortening strain at lower activations. We attempted this for one preparation, stimulating the muscle supra- and submaximally at both 4 and 6 Hz oscillation frequencies. We found that total fascicle strain during submaximal stimulation (*P*_30_) decreased from 8 to 5.8% *L*_0_ (i.e., a 27.2 % reduction) while fascicle strain during supramaximal stimulation decreased from 9 to 7.5% *L*_0_ (i.e., a 17% reduction). While not definitive, this single observation is consistent with the hypothesis that inactive muscle tissue mass drives the reduction in strain at submaximal activations.

An alternative, and not mutually-exclusive explanation for reduced strain at lower activation is interactions between force–length effects and series elastic compliance rather than inertia *per se*. With decreased activation, relative load approaches a larger fraction of the instantaneous force capacity of the muscle, limiting both shortening velocity and excursion as predicted by classic after-load force-velocity experiments (Hill, 1938; Katz, 1939). In our preparation, the lever motor imposed a fixed sinusoidal length trajectory on the muscle, in which additional fascicle shortening had to be accommodated by stretching series elastic elements or increasing force on the lever. Despite efforts to minimize series-elastic compliance, substantial fixed-end compliance remained with fascicles shortening approximating 8% *L*_o_ during isometric tetani, and oscillatory fascicle strain (±5.7%) exceeding imposed muscle strain (±3% *L*_o_). At lower levels of activation, the force available to stretch series elastic elements is reduced, limiting fascicle shortening and increasing apparent lengthening during the imposed cycle.

### Effects of submaximal stimulation on MTU-fascicle latency

We observed significant effects of stimulation intensity on the timing of muscle tendon unit (motor) lengthening and shortening versus fascicle lengthening and shortening. With respect to lengthening latency, the model-predicted effect amounted to an overall lag of approximately 5.8 ms, or 2.3% of the entire work loop cycle (at 4Hz) for *P*_30_, as compared to the timing of fascicle length change at *P*_0_. The effects we observed in the timing of length reversal (lengthening to shortening and the reverse) suggest two disparate, yet related effects of inactive muscle mass: one of acceleration and the other of deceleration. As the motor reaches peak length, and then begins to drive muscle shortening, the muscle fascicles must accelerate to a shortening velocity greater than the motor in order to produce force. At supramaximal stimulation, the muscle fascicles increase shortening velocity more rapidly relative to submaximal activation. Although we did not measure motor or fascicle velocity directly (as motor velocity – and thus velocity of the MTU – constantly changes during sinusoidal work-loops), we found that fascicle shortening occurred significantly later than the motor during submaximal contractions versus during maximal contractions (Figure 6). This pattern is consistent with previous findings for the rat PL and other small muscles during isotonic shortening (Holt et al., 2014; Tijs et al., 2021) showing maximal shortening velocity decreasing from 3.52 to 1.87 *L*_0_s^-1^ between maximal and 30% activation.

Although fascicle lengthening latency could be a consequence of force-length effects as discussed above, it is unclear how this could account for latency differences for fascicle deceleration. As the motor reverses from shortening to lengthening, the fascicles must decelerate their rate of shortening prior to being passively lengthened. We found that during submaximal stimulation, the fascicles continued to shorten for a significantly greater duration, into motor lengthening, than during maximal stimulation (Figure 6). In other words, at submaximal stimulation, there is a greater temporal decoupling in the behavior between the motor—and therefore muscle tendon unit—and the underlying fascicles. In the absence of an effect of inactive muscle tissue mass, one would expect that at lower activation, the muscle is less able to generate sufficient force to stretch compliant elements and shorten against the lengthening muscle tendon unit. Instead, it is likely that the inactive mass causes continued passive fascicle shortening, even after stimulation has ceased. We contend that the increased lag at the length reversals between shortening and lengthening are likely caused by the added inactive tissue mass that causes a reduction in fascicle length change acceleration and deceleration during submaximal contractions. Prolonged shortening, as a result, may have increased positive work (whether the fascicles were active or not), and compensated for work reductions during passive fascicle acceleration.

### Study limitations and future directions

Our study has several limitations. Using rat PL, a small and mixed-fibered, though predominantly fast muscle limits the generalizability of our findings to larger muscles of the sizes found in humans. However, our approach is a logical first step that provides an initial and likely conservative estimate of the size limit for inertial effects to manifest. Indeed, as per scaling principles of mass versus force-dependent fiber cross-sectional area, inertial effects are expected to increase with muscle and organism size. For now, our testing of a size-dependent hypothesis in a small muscle system may bias towards an underestimation of inertial penalties that arise from inactive muscle mass.

Despite best efforts, our reductionist approach involving surgical isolation compromised the naturally homeostatic muscle environment, and our artificial stimulation approach is unlikely to replicate organismal neuromuscular control. More importantly, oscillating at a fixed frequency with a sinusoidal trajectory may suppress inertial effects that otherwise manifest at faster movement and more abrupt motion transitions between power- and recovery strokes in natural muscle-based movement. Although our preparation removed as much series elastic compliance from the muscle preparations as possible, the remaining aponeurosis likely masks inertia-driven alterations in fascicle contractile dynamics, and averaging our data across individuals, muscle regions, and work loop cycles may also obscure small, spatially or temporally localized effects.

Combined, these factors may collectively mask subtle inertial contributions to muscle performance that in the future can be better understood by (a) testing larger muscles with and without pennate architecture, and (b) by using our empirical data from isolated muscles in computational approaches to determine how an expanded input parameter-set, with varying oscillation frequencies, strain-ranges, and stimulation phasing, may amplify or attenuate fluctuations in inertial effects from submaximal muscle contractions.

## Conclusions

Whereas our results suggest some effects of inertia at this small muscle size, specifically on the temporal dynamics and extent of fascicle contractile cycling strain, these effects were not substantial enough to significantly influence muscle work production. However, given that our experimental design might mask subtle as well as transient effects of inertia, and a similar study found more substantial evidence at comparable muscle size, we predict that future work on larger muscles will recover meaningful inertial effects at muscle sizes more relevant to human movement.

## Acknowledgements

We thank Pedro Ramirez for animal care.

## Funding

Research reported in this publication was supported by NIAMS under NIH Award Number R01AR080797.

## Disclosures/ competing interests

Authors have nothing to disclose and no competing interests

## Data and code availability

All data collected and code required to replicate the main findings of the paper are available upon request

## Notes

### Competing Interest Statement

The authors have declared no competing interest.

